# Rostral Anterior Cingulate Activations inversely relate to Reward Payoff Maximation & predict Depressed Mood

**DOI:** 10.1101/2021.06.11.447974

**Authors:** Pragathi Priyadharsini Balasubramani, Juan Diaz-Delgado, Gillian Grennan, Fahad Alim, Mariam Zafar-Khan, Vojislav Maric, Dhakshin Ramanathan, Jyoti Mishra

**Affiliations:** Neural Engineering and Translation Labs, Department of Psychiatry, University of California, San Diego, La Jolla, CA, USA; Department of Mental Health, VA San Diego Medical Center, San Diego, CA

**Keywords:** reward prediction, risk sensitivity, reinforcement learning, anterior cingulate cortex, depression

## Abstract

Choice selection strategies and decision making are typically investigated using multiple-choice gambling paradigms that require participants to maximize reward payoff. However, research shows that performance in such paradigms suffers from individual biases towards the frequency of gains to choose smaller local gains over larger longer term gain, also referred to as melioration. Here, we developed a simple two-choice reward task, implemented in 186 healthy human adult subjects across the adult lifespan to understand the behavioral, computational, and neural bases of payoff maximization versus melioration. The observed reward choice behavior on this task was best explained by a reinforcement learning model of differential future reward prediction. Simultaneously recorded and source-localized electroencephalography (EEG) showed that diminished theta-band activations in the right rostral anterior cingulate cortex (rACC) correspond to greater reward payoff maximization, specifically during the presentation of cumulative reward information at the end of each task trial. Notably, these activations (greater rACC theta) predicted depressed mood symptoms, thereby showcasing a reward processing marker of potential clinical utility.

**Significance Statement:** This study presents cognitive, computational and neural (EEG-based) analyses of a rapid reward-based decision-making task. The research has the following three highlights. 1) It teases apart two core aspects of reward processing, i.e. long term expected value maximization versus immediate gain frequency melioration based choice behavior. 2) It models reinforcement learning based behavioral differences between individuals showing that observed performance is best explained by differential extents of reward prediction. 3) It investigates neural correlates in 186 healthy human subjects across the adult lifespan, revealing specific theta band cortical source activations in right rostral anterior cingulate as correlates for maximization that further predict depressed mood across subjects.

## Introduction

Cognitive and neural responses to reward and risk are quintessential to understanding human behavior. Gambling tasks predominantly form the experimental test beds for measuring reward and risk processing abilities in humans (Bechara et al., 1997; Brevers et al., 2013). However, it is widely debated how well these tasks can separate decision-making based on frequency of gains/losses versus expected value of payoff for different choice-sets (Bechara et al., 1997, 2000, 2000; Bembich et al., 2014; Christakou et al., 2009; Garon et al., 2006; R. Gupta et al., 2011; Mueller et al., 2010; Must et al., 2013; Oberg et al., 2011; Roca et al., 2008; Woodrow et al., 2019).

For instance, studies that have controlled for gain frequency in the Iowa Gambling Task (IGT) show that subject choices reflect their inherent gain frequency preferences in the task, which portray relatively immediate reinforcement based choice behavior rather than a behavior that maximizes expected value or long-term payoff (Chiu et al., 2008; Chiu & Lin, 2007; Lin et al., 2009; Napoli & Fum, 2010; Singh & Khan, 2012). In contrast, some researchers suggest such local minima choices also referred to as melioration may be the rational solution in the face of uncertainty (Sims et al., 2013). Additionally, task related performance measures for long-term payoff versus immediate gain frequency bias may reflect distinct reinforcement learning and decision making strategies, e.g. sensitivity to rewards and risks, learning rate, and other behavioral execution strategies (Franken & Muris, 2005; Furl, 2010; Harman, 2011; Newman et al., 2008; Weller et al., 2010), which prior neural studies of reward payoff maximization have not accounted for.

In this study, we uniquely separate immediate gain frequency bias driven decision-making from advantageous longer term payoff-based decision making. Specifically, we designed a two-choice paradigm with two distinct blocks – a Δ_0_payoff (baseline) block where two reward choice options have equal payoffs and reward variance suitable for measuring the immediate gain frequency bias, and a Δpayoff (difference) block where the two-choice options have unequal payoffs suitable for measuring payoff influences. We thereby, tease apart measurements of immediate gain frequency biased response from expected value or long-term payoff based response, to understand the distinct cognitive and neural mechanisms underlying payoff decisions.

Second, we capitalize on computational reinforcement learning (RL) models to understand the basis of individual differences in reward and risk based learning across subjects (Balasubramani et al., 2014; Balasubramani & Chakravarthy, 2019; Chakravarthy et al., 2018; A. Gupta et al., 2013; Muralidharan et al., 2014). The RL framework provides the ability to simulate the observed behavior and estimate the hidden parameters forming the basis for individual differences in learning and performance. Specifically in this study, we checked whether the observed subject behavior for reward payoff maximization versus gain frequency bias melioration can be explained by extent of future reward prediction (Doya, 2002), or differential risk seeking towards gains versus losses (Balasubramani et al., 2014).

Neurally, earlier studies have suggested the significant role of frontal executive regions, particularly the medial prefrontal cortex (mPFC) in reward and risk based performance (Balasubramani & Hayden, 2018; Bechara et al., 1997, 2000; Brevers et al., 2013; Kennerley et al., 2011; Moccia et al., 2017). Uniquely, in this study, we estimate the neural correlates for payoff relevant decisions while accounting for significant RL parameters and individual differences in immediate gain frequency biases. Finally, we understand the relationship between the identified neural correlates for reward maximization to individual’s self-reported mood symptom variations. We show that reward payoff maximization correlates within mPFC are sensitive to depressed mood, and hence our experimental and computational framework for identifying the regions of interest in mPFC may serve future clinical utility as a biomarker for depression.

## Methods

### Participants

198 adult human subjects (age mean ± standard deviation 35.44 ± 20.30 years, range 18-80 years, 115 females) participated in the study. All participants provided written informed consent for the study protocol approved by the University of California San Diego institutional review board (UCSD IRB #180140). Twelve of these participants were excluded from the study as they had a current diagnosis for a psychiatric disorder and current/recent history of psychotropic medications for a final sample of 186 healthy adult participants. All participants reported normal/corrected-to-normal vision and hearing and no participant reported color blindness. For older adults >60 years of age, participants were confirmed to have a Mini-Mental State Examination (MMSE) score >26 to verify absence of apparent cognitive impairment (Arevalo-Rodriguez et al. 2015). All data was collected prior to the COVID-19 period of restricted human subjects research.

### Surveys

All participants provided demographic information by self-report including age, gender, race (in a scale of 1 to 7: Caucasian; Black/African American; Native Hawaiian / Other Pacific Islander; Asian; American Indian / Alaska Native; More than one race; Unknown or not reported) and ethnicity; socio-economic status (SES) was measured on the Family Affluence Scale from 1 to 9 (Boudreau and Poulin, 2008), and any current/past history of clinical diagnoses and medications were reported. For older adults >60 years of age, participants completed the Mini-Mental State Examination (MMSE) and scored >26 to verify absence of apparent cognitive impairment (Arevalo-Rodriguez et al., 2015). All participants completed subjective mental health self-reports using standard instruments, ratings of inattention and hyperactivity obtained on the Adult ADHD Rating Scale (New York University and Massachusetts General Hospital. Adult ADHD-RS-IV with Adult Prompts. 2003; : 9–10), Generalized Anxiety Disorder 7-item scale GAD-7 (Spitzer et al., 2006) and depression symptoms reported on the 9-item Patient Health Questionnaire, PHQ-9 (Kroenke et al., 2001). Symptoms for these psychiatric conditions were measured because they have been related to changes in reward processing (Admon & Pizzagalli, 2015; Dillon et al., 2014; Luman et al., 2010).

### Task Design

We investigated a two-choice decision-making task that enabled a rapid assessment and was easy to understand across the adult lifespan. In this task that we refer to as *Lucky Door*, participants chose between one of two doors, either a rare gain door (RareG, probability for gains p=0.3, for losses p=0.7) or a rare loss door (RareL, probability for losses p=0.3, for gains p=0.7). Participants used the left and right arrow keys on the keyboard to make their door choice. Door choice was monitored throughout the task. In two separate blocks, we investigated whether the overall expected value (payoff) of the choice door can influence individual behavior. In the baseline block with Δ_0_payoff (no-payoff difference), the two choice doors did not differ in payoff (RareL door, p=0.3 for -70 coins and p=0.7 for +30 coins, payoff=0; RareG door, p=0.3 for +70 coins and p=0.7 for -30 coins, payoff=0). In the experimental difference block with Δpayoff (payoff difference), expected value or payoff was greater for the RareG door (p=0.3 for +60 coins, p=0.7 for -20 coins, payoff=+40) than for the RareL door (p=0.3 for -60 coins, p=0.7 for +20 coins; payoff=-40). Manipulation of payoff, with greater expected value tied to the RareG door, allowed for investigating individual propensities to prioritize long-term (or cumulative) vs. short-term (or immediate) rewards. The RareG door was assigned greater payoff because selecting this door suggests payoff-based decision processing in subjects as opposed to simply choosing based on frequency of gains, in which case the RareL choice should be preferred. 40 trials were presented per block and block order was randomized across participants; two practice trials preceded the main Δpayoff or Δ_0_payoff blocks. **Figure 1A** shows a schematic of the task stimulus sequence and **Supplementary table 1** shows the reward distribution that was shuffled and updated after every 10 trials had been sampled from that set.

**Figure 1.**
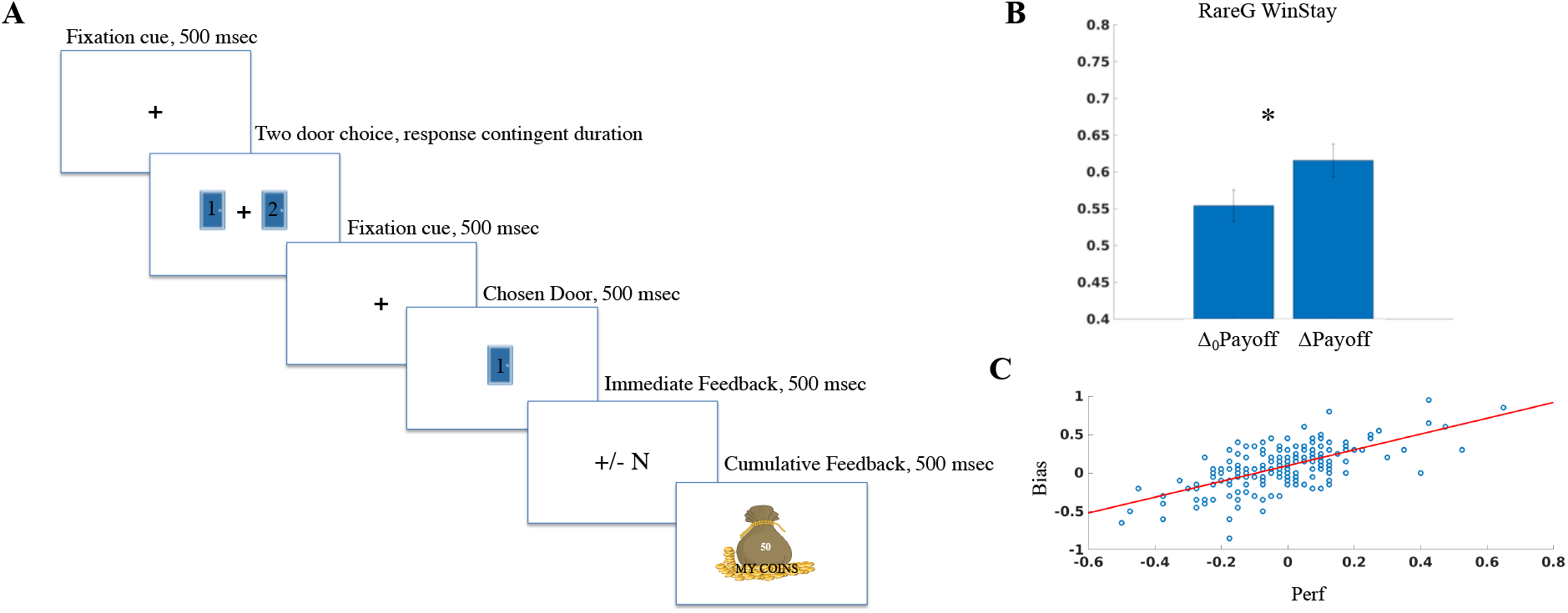
Reward task and associated behavior. **A)** As per the task schematic, participants fixated for 0.5 sec, then chose from one of two choice doors. Post-response, fixation reappeared for 0.5 sec, followed by presentation of the chosen door for 0.5 sec, then immediate gain or loss feedback provided for 0.5 sec, and finally, cumulative feedback of all gains/losses up to the present trial shown for 0.5 sec. Reward distributions for the door choices are presented in Supplementary Table 1. **B)** Win-Stay behavior for the rare gain RareG door is significantly greater on the ΔPayoff versus Δ_0_Payoff block. *:p<.05. **C)** Gain frequency *Bias* significantly predicts payoff performance, *Perf* (r=0.65, p<0.0001).

The *Lucky Door* task was deployed in Unity as part of the assessment suite on the *BrainE* (short for Brain Engagement) platform (Balasubramani et al., 2020). The Lab Streaming Layer (LSL (Kothe et al., 2019)) protocol was used to time-stamp each stimulus/response event during the task. Study participants engaged with the assessment on a Windows 10 laptop sitting at a comfortable viewing distance.

### Electroencephalography (EEG)

EEG data was collected simultaneous to the *Lucky Door* task using a 24-channel Smarting device with a semi-dry and wireless electrode layout (Next EEG— new human interface, MBT). Data were acquired at 500 Hz sampling frequency at 24-bit resolution. Cognitive event markers were integrated using LSL and data files were stored in xdf format.

### Behavioral analyses

Task speeds were calculated as log(1/RT), where RT is response time in seconds. We computed the payoff sensitive performance response (*Perf*) as the difference in proportion selection of the RareG door between the Δpayoff and the Δ_0_payoff blocks; RareG vs. RareL EVs differed only in the Δpayoff block. We computed gain frequency bias (*Bias*) as the difference in selected proportion of RareL and RareG doors in the Δ_0_payoff block where the payoff for both the doors was the same. While *Perf* is indicative of subjective payoff maximization based selection of advantageous choices, *Bias* informs the choice bias and melioration for higher immediate gain frequencies. For N fraction of responses in each block, we calculated:

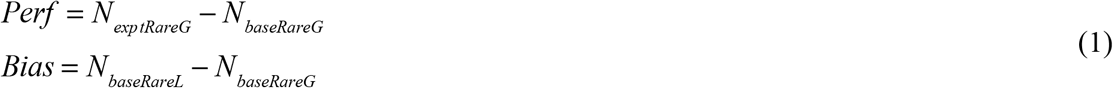

We also calculated Win-Stay and Lose-Shift performance on both Δpayoff and Δ_0_payoff blocks. Win-Stay was computed as the proportion of times the subject repeated the same choice option in the next trial after obtaining a gain for choosing that option in the current trial. Lose-Shift was computed as the proportion of times the subject would shift away from the current choice option in the next trial, on obtaining a loss in the current trial.

### Reinforcement Learning (RL) Model

We simulated three parsimonious RL models with up to two free parameters, which have a key role in reward-based decision-making, to explain the observed individual subject behavior; see **Figure 2A** for model schematic (Balasubramani et al., 2014; Sutton & Barto, 1998):

1. Model A optimized the time scale of reward prediction (γ) for every subject;
2. Model B optimized risk sensitivity (α) for every subject; and
3. Model C optimized both γ and α to for every subject.

**Figure 2.**
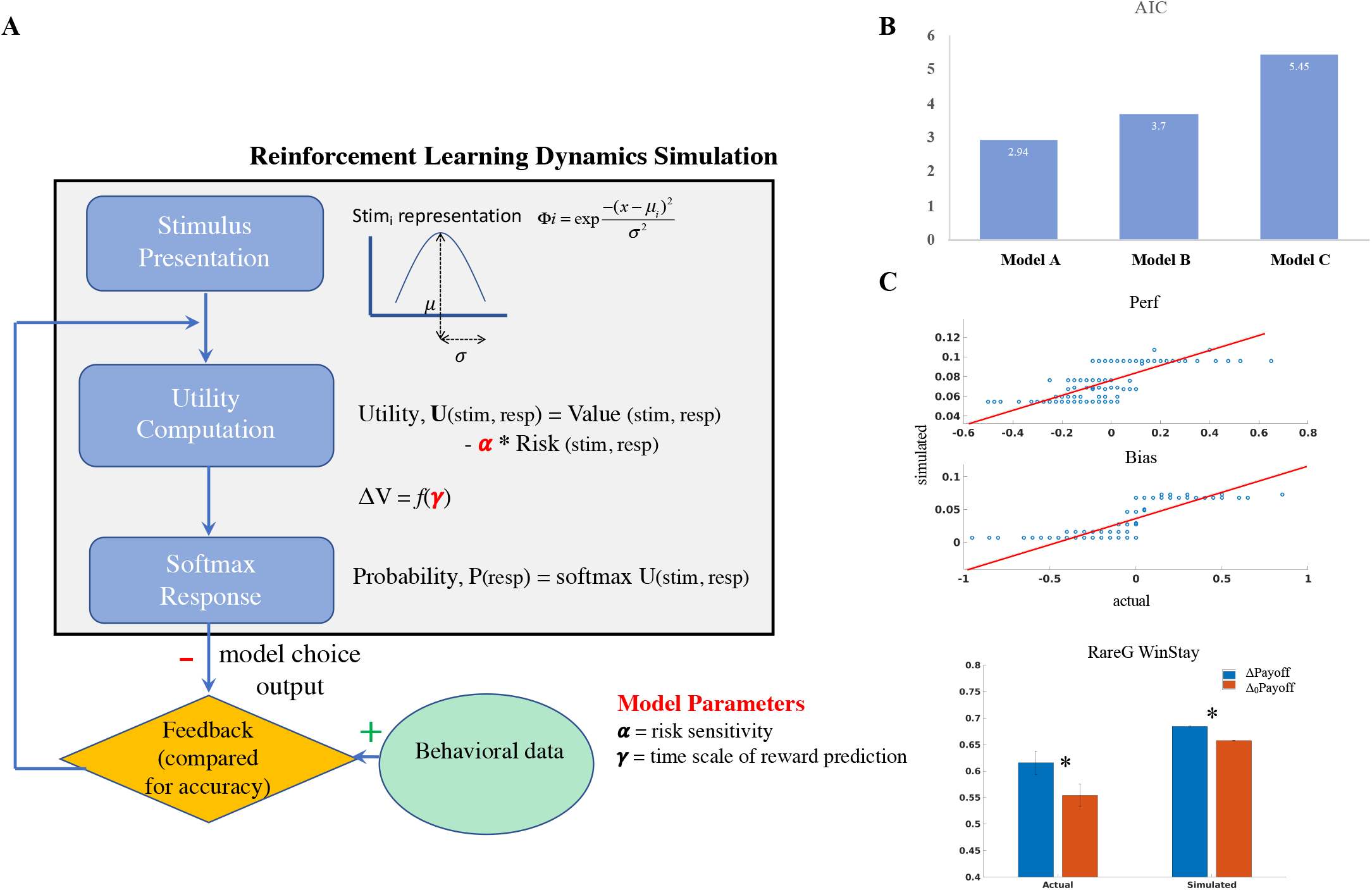
Reinforcement Learning Model. **(A)** Schematic representing the stimulus, value function and choice selection modules. The model results for number of selections associated with each of the choice door stimuli in each task block are compared against the actual selections made by each subject, for purposes of model optimization. The model uses the utility, U, associated with each choice response for making the decision, where the utility is a function of reward average and reward variance associated with choices. The decision in the model is taken using the SoftMax probability, P, of making the choices. Model parameters are highlighted as α (model agent’s differential risk sensitivity to gain and loss outcome uncertainties), and γ (time scale of reward prediction). **(B)** AIC values for three models, Model A (γ optimized), B (α optimized), C (γ and α optimized), show that the γ model was best performing. **(C)** The γ optimized model showed strong correlations between simulated and actual *Perf* (ρ(185)=0.81, p<0.0001), *Bias* (ρ(185)=0.89, p<0.0001) and RareG Win-Stay difference between blocks (ρ(185)=0.17 p=0.004).

The time scale of reward prediction parameter (γ) represents whether reward prediction is myopic or long-sighted, lower values γ ∈ (0 1), γ → 0 suggest myopic reward prediction leading to impulsive decisions while higher values, γ → 1 suggest long-sighted integration of rewards for decisions.

The risk sensitivity parameter (α) measures the extent to which expected uncertainty associated with the door influences the decision utility, the smaller the parameter value α ∈ (−1 1), α → -1 the higher is risk seeking, while a larger value, α → 1 indicates high risk aversiveness.

The simulation agent had reward distributions as in the real experiment but scaled down by multiplying with a parameter 0.1, and varying with blocks (Δpayoff, Δ_0_payoff) that were randomly ordered. There were as high as 50,000 trials in each block for letting model performance converge.

The agent has to choose between two doors each of which (stimulus, *s*) was represented by a radial basis function (Φi) as below:

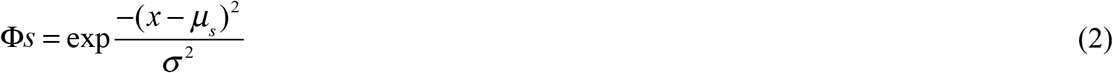

Here, the μ_s_ and σ denotes the mean (*s* ∈ [1 2]; door1 = 1; door2 = 2) and standard deviation of the inverse attention parameter, respectively. σ is set to 1 in our models.

The door stimulus is multiplied with the weight matrix *wv* for computing its value function, *Q*, and *wr* for constructing its risk function, *√h*.

Utility associated with any state at a trial, *t*, is the combination of value and risk function(Bell, 1995; d’Acremont et al., 2009), where the risk function is modulated by a risk sensitivity parameter α. Higher the α, higher the risk aversiveness of the subject.

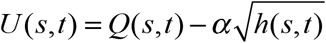

Where

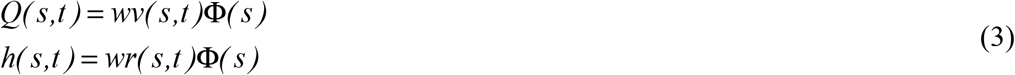

The door choice selection is performed using the SoftMax principle defined as below. According to SoftMax, the probability for choosing a door at trial, *t*, is *P*(s,t):

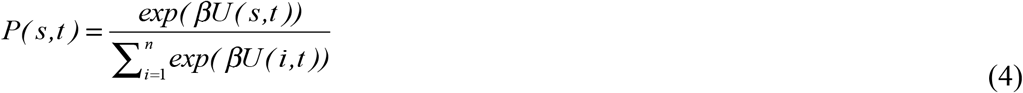

Here, *n* is the total number of doors available, and *β* is the exploration index. β is set to 1 in our models.

After choice selection, the weight functions are updated using the below principles. The choice value function *Q* at trial *t*+1 for door, *s*, may be expressed as,

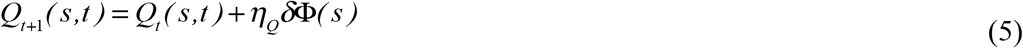

where *ηQ* is the learning rate of the value function (0 < *ηQ* < 1) for the stimulus variable, Φ(s). δ is the temporal difference error represented as

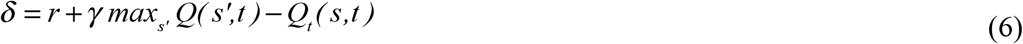

where *r* is the reward associated with taking an action, a, for stimulus, s, at time, t, and γ is the time scale of reward prediction. Similar to the value function, the risk function h has an incremental update as defined by the below equation. Optimizing the risk function in addition to the value function is shown to capture human behavior well in a variety of cognitive tasks involving rewards, punishments and risk (Balasubramani, Chakravarthy, Ravindran, et al., 2015; Balasubramani et al., 2014).

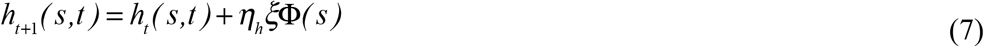

where *ηh* is the learning rate of the risk function (0 <*ηh*< 1), and *ξ* is the risk prediction error expressed by the below equation.

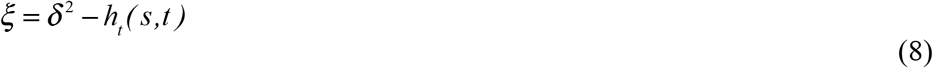

For simplicity, we model as *η*_*h*_ = *η*_*Q*_ =0.1 as an initial optimization for our subjects for η provided a median of 0.1. The weights *wv* and *wr* are set to a small random number from set [-0.0005 0.0005] at trial = 1. The weights are normalized by dividing by their norm.

The cost function optimizes the frequency of selections of rare gain and rare loss options in Δ_0_payoff and Δpayoff blocks for every subject after running the simulation agent for 10 instances of 100,000 each, and inferring the optimal parameters for every participant in our study using *fmincon* function in MATLAB. Cost function = sum of squares of the difference for observed actual (Proportion# RareG_expt_ + Proportion# RareL_expt_ + Proportion# RareG_base_ + Proportion# RareL_base_) – simulated actual (Proportion# RareG_expt_ + Proportion# RareL_expt_ + Proportion# RareG_base_ + Proportion# RareL_base_). Optimization is carried out for either one (Models A,B) or two (Model C) parameters, using *fmincon*(). We ran fmincon() 100 times to choose the parameter set with least cost for any subject.

The AIC (Akaike Information Criteria) for these models were built using the likelihood function, which was estimated as the average correlation coefficient between the simulated and the observed key task behaviors - 1) payoff performance or *Perf* measure (eqn. 1); 2) gain frequency *Bias* measure (eqn. 1); and 3) the block differences between Δpayoff and Δ0payoff blocks in Win-Stay results for the RareG door.

### Neural data processing

We applied a uniform processing pipeline to all EEG data acquired simultaneous to the reward task. This included: 1) data pre-processing, 2) computing event related spectral perturbations (ERSP) for all channels, and 3) cortical source localization of the EEG data filtered within relevant theta, alpha and beta frequency bands.

1. Data preprocessing was conducted using the EEGLAB toolbox in MATLAB(Delorme & Makeig, 2004). EEG data was resampled at 250 Hz, and filtered in the 1-45 Hz range to exclude ultraslow DC drifts at <1Hz and high-frequency noise produced by muscle movements and external electrical sources at >45Hz. EEG data were average referenced and epoched to the chosen door presentation during the task, in the -.5 sec to +1.5 sec time window (**Figure 1**). Any missing channel data (one channel each in 6 participants) was spherically interpolated to nearest neighbors. Epoched data were cleaned using the autorej function in EEGLAB to remove noisy trials (>5sd outliers rejected over max 8 iterations; 0.91± 2.65% of trials rejected per participant). EEG data were further cleaned by excluding signals estimated to be originating from non-brain sources, such as electrooculographic, electromyographic or unknown sources, using the Sparse Bayesian learning (SBL) algorithm(Ojeda et al., 2018, 2021), https://github.com/aojeda/PEB) explained below in the cortical source localization section.
2. For ERSP calculations, we performed time-frequency decomposition of the epoched data using the continuous wavelet transform (cwt) function in MATLAB’s signal processing toolbox. Baseline time-frequency (TF) data in the -250 ms to -50 ms time window prior to chosen door presentation were subtracted from the epoched trials (at each frequency) to observe the event-related synchronization (ERS) and event-related desynchronization (ERD) modulations (Pfurtscheller, 1999)..
3. Cortical source localization was performed to map the underlying neural source activations for the ERSPs using the block-Sparse Bayesian learning (SBL) algorithm (Ojeda et al., 2018, 2021) implemented in a recursive fashion. This is a two-step algorithm in which the first-step is equivalent to low-resolution electromagnetic tomography (LORETA) (Pascual-Marqui et al., 1994). LORETA estimates sources subject to smoothness constraints, i.e. nearby sources tend to be co-activated, which may produce source estimates with a high number of false positives that are not biologically plausible. To guard against this, SBL applies sparsity constraints in the second step wherein blocks of irrelevant sources are pruned. This data-driven sparsity constraint of the SBL method reduces the effective number of sources considered at any given time as a solution, thereby reducing the ill-posed nature of the inverse mapping. In other words, one can either increase the number of channels used to solve the ill-posed inverse problem or impose more aggressive constraints on the solution to converge on the source model. The two-stage SBL produces evidence-optimized inverse source models at 0.95AUC relative to the ground truth while without the second stage <0.9AUC is obtained (Ojeda et al., 2018, 2021). Prior research has also shown that sparse source imaging constraints can be soundly applied to low channel density data (Ding & He, 2008; Stopczynski et al., 2014). Furthermore in Balasubramani et al. (42) we have shown that the ROI estimates resulting from this cortical source mapping have high test-retest reliability (Cronbach’s alpha = 0.77, p<0.0001).

Source space activity signals were estimated and their root mean squares were partitioned into (1) regions of interest (ROIs) based on the standard 68 brain region Desikan-Killiany atlas (Desikan et al., 2006), using the Colin-27 head model (Holmes et al., 1998) and (2) artifact sources contributing to EEG noise from non-brain sources such as electrooculographic, electromyographic or unknown sources; activations from non-brain sources were removed to clean the EEG data. Cleaned subject-wise trial-averaged EEG data were then specifically filtered in theta (3-7 Hz), alpha (8-12 Hz), and beta (13-30 Hz) bands and separately source localized in each of these bands to estimate their cortical ROI source signals. The source signal envelopes were computed in MatLab (envelop function) by a spline interpolation over the local maxima separated by at least one time sample; we used this *spectral amplitude* signal for all neural analyses presented here. For analyses, we focused on theta, alpha and beta signals in the relevant selected choice period (0-500 ms after selected choice presentation), trial reward period (500-1000 ms after selected choice presentation), and the cumulative reward period (1000-1500 ms after selected choice presentation).

### Statistical Analyses

We fit robust multivariate linear regression models in MATLAB to investigate the behavioral relationships between the *Perf* measure and demographic variables (age, sex, race, ethnicity and SES), while controlling for *Bias* and order of block presentation. For any linear regression model, the response variable was log-transformed for normality and we identified significant factors contributing to the main effects. For regression models, we report the overall model R^2^ and p-value, and individual variable *β* coefficients, t-statistic, degrees of freedom, and p-values.

Channel-wise theta, alpha, beta ERS and ERD modulations for significant spectral activity were computed relative to baseline by first processing for any outliers; any activations greater than 5MAD from the median were removed from further analyses. The significant average activity across all trials were found by performing t-tests (p<0.05) across subjects, followed by false discovery rate (FDR, alpha = 0.05) corrections applied across the three dimensions of time, frequency, and channels (Genovese et al., 2002).

For computing source level activity correlates of the behavioral *Perf* measure, we first found the difference in RareG door specific neural activations between Δpayoff and Δ_0_payoff blocks in three frequency bands – theta, alpha and beta and in three relevant trial periods: selected choice presentation, trial reward and cumulative reward period. We again used robust linear regression fits for identifying individual ROIs that relate to the *Perf* measure, while also accounting for individual *Bias* and variations in RL as predicted by the best fitting RL model. The results were family-wise error rate corrected for multiple comparisons for 3 trial periods and 3 frequency bands (FWER correction, p<0.0055). The independently identified ROIs were further factored in a unified multivariate linear regression model to account for comparisons across ROIs; significant ROIs in this final multivariate model were reported after controlling for *Bias*, and the RL model-fit parameter (p<0.05).

Finally, we used robust multivariate linear regressions to model the self-reported mood symptoms of Anxiety, Depression, Inattention, Hyperactivity using predictors of demographic variables, *Perf, Bias*, RL model-fit parameter along with the identified neural correlates of *Perf*. Adjusted responses from robust multivariate models were plotted using the plotAdjustedResponse function in MATLAB.

## Results

### Options associated with larger payoff show win-stay behavior and their choice fraction is proportional to individual gain frequency bias

Healthy adult subjects (N = 186, ages 18-80 years, 115 females) performed a two-choice gambling task, *Lucky Door*, which implemented two distinct blocks of choices; the Δ_0_payoff block delivered choice-sets with different gain frequencies but no differences in payoff, while the Δpayoff block delivered choice-sets with same gain frequencies as the Δ_0_payoff block yet with long-term payoff (i.e. expected value) differences (**Figure 1A**; **Supplementary Table 1**). Specifically, the Δ_0_payoff block only varied the gain frequency associated with the choice doors, with one door leading to 70% positive reward outcomes (Rare Loss or RareL door) while the other resulting in 70% negative reward outcomes (Rare Gain or RareG door), yet maintaining the same reward average or long-term expected value/payoff. The Δpayoff block, presented in a random sequence order relative to the Δ_0_payoff block across subjects, had the same gain and loss frequency setup as the Δ_0_payoff block, but the rewards associated with the RareG door resulted in a larger long-term payoff than the RareL door. Participants executed 40 trials per block. Thus, we calculated the payoff related performance measure, *Perf* as the difference in probability of RareG selections on Δpayoff vs. Δ_0_payoff blocks. Gain frequency *Bias* was measured as the difference in proportion of RareL vs. RareG choices on the Δ_0_payoff block.

In terms of subject behavior, the actual proportion of RareG choices between the two blocks did not differ (paired Wilcoxon signed-rank test: N_RareG_ Δpayoff:17.74±0.39; Δ_0_payoff:18.47±0.40; p=0.07). But, we found that Win-Stay behavior for the RareG choices distinguished performance on the two blocks (mean±SEM Win-Stay RareG, Δpayoff:0.61±0.02; Δ_0_payoff:0.55±0.02; block difference: 0.06± 0.03, p=0.02, **Figure 1B**). Corresponding lose-shift behavior did not differ between blocks (p=0.33).

Additionally, we found that the payoff-based responses, *Perf* were significantly correlated to individual gain frequency *Bias* (r=0.65, p<0.0001, **Figure 1C**).

Next, we implemented multivariate regression to model the payoff-related performance, *Perf*, based on all self-reported demographic (age, gender, race, ethnicity, socio-economic status SES) and mental health (anxiety, depression, inattention and hyperactivity) predictors as per **Table 1**. This model also included individual gain frequency *Bias* and order of block presentation. The overall *Perf* model was significant (R^2^=0.43, p<0.0001). Interestingly the only variable that significantly predicted *Perf* was Gain frequency *Bias* (β=0.37 ±0.04, t(151)=9.18, p<0.0001). Thus, hereafter, we control for *Bias* in all payoff-relevant analyses below.

**Table 1.** Subject characteristics. Median ± MAD for subjects demographics variables, mental health self-report scores, and parameter γ from the preferred reinforcement learning model. MAD: median absolute deviation, SES: socioeconomic status score.

| Demographics | | Median ± MAD |
| --- | --- | --- |
| Age | | 25.00 ± 14.87 |
| Gender n (%) |  |  |
|  | Male | 71 (38.17) |
|  | Female | 115 (61.83) |
| Ethnicity n (%) |  |  |
|  | Caucasian | 116 (62.37) |
|  | Black or African American | 4 (2.15) |
|  | Native Hawaiian, Pacific Islander | 0 (0) |
|  | Asian | 37 (19.89) |
|  | American Indian, Alaska Native | 4 (2.15) |
|  | Multi-racial | 12 (6.45) |
|  | Others | 12 (6.45) |
| Race n (%) |  |  |
|  | Hispanic or Latino | 25 (13.51) |
|  | Not Hispanic or Latino | 155 (83.78) |
|  | Unknown | 5 (2.70) |
| SES | | 5.00 ± 1.34 |
| Mental Health | | Median ± MAD |
| Anxiety | | 3 ± 2.88 |
| Depression | | 3 ± 2.76 |
| Inattention | | 4 ± 3.99 |
| Hyperactivity | | 3 ± 2.96 |
| Behavior | | Median ± MAD |
| Perf | | -0.02±0.13 |
| Bias | | 0.10±0.21 |
| Model $\gamma$ | | 0.99± 0.07 |

### Individual differences in reward prediction explain task behavior

Prior RL modeling studies have suggested that task behavior can be explained with greater accuracy when both the long term expected value or payoff and the expected risk are accounted for during decision-making (Balasubramani, Chakravarthy, Ravindran, et al., 2015; Balasubramani et al., 2014). The major advantages of building these RL models are that the models can characterize the individual differences in each participant in terms of computational parameters that manifest in learning and executive control. Secondly, they can explain the converged behavioral dynamics in each participant sans experimental trial limitations.

Using this RL framework, we were interested in investigating how are choice decisions affected by 1) extent of integration of rewards over time i.e. time scale of reward prediction, and 2) differential risk sensitivity to gains and losses affecting the choices, where risk is the variance in reward outcomes. In order to find whether the observed subjective behavioral differences are driven by one of the two or both of the above decision making measures, we built three separate reinforcement learning models (RL, **Figure 2A**) that simulate behavior by optimizing time scale of reward prediction (γ: model A), risk sensitivity (α: model B) or both γ and α parameters (model C). In model A, higher γ parameter values represent long-sightedness; risk sensitivity α is set to 0 in this model. In model B, higher α parameter values indicate risk aversiveness and lower values indicate risk seeking; the reward prediction γ parameter is set to 0 in this model. All models were optimized using robust solvers in MATLAB (see Methods to find global minima with multiple random initial conditions) to match to the observed choice selection behavior observed in each block.

Next, we computed the likelihood of any Model A, B, C to explain the significantly observed behavioral measures: which are *Perf, Bias* and the block differences in proportion of *Win-Stay* to the RareG door. The likelihood, taken as correlations between the simulated and actually observed, was used to construct model AIC (Akaike Information Criterion). The AIC values for Models A, B and C were 2.94, 3.70 and 5.45, respectively (**Figure 2B**), suggesting that Model A can preferably explain the observed behavior (**Figure 2C**). This means the individual differences in the time scale of reward prediction can preferably explain the reward maximization (*Perf*) as well as melioration (*Bias*) behavior. On parameter recovery analysis for this model, the simulated model solutions were significantly correlated with the recovered solutions, implying that our model can reliably explain the subjective behaviors (Wilson & Collins, 2019).

### Right rostral anterior cingulate cortex encodes reward payoff maximization

Participants performed the reward task with simultaneous EEG, which we analyzed in the theta (3-7 Hz), alpha (8-12 Hz), and beta (13-30 Hz) frequency bands in cortical source space parcellated as per the Desikan-Killiany regions of interest (Desikan et al., 2006). To identify the neural correlates underlying reward payoff maximization (*Perf*), we modeled the neural variables as predictors of *Perf* using robust multivariate linear regression, while accounting for gain frequency *Bias* that was significantly related to *Perf* (**Figure 1**), and the optimal RL model parameter (γ).

We investigated neural activations from three relevant trial periods: immediately post-presentation of selected choice but prior to reward (0-500 ms selected choice period), during presentation of trial reward (0-500 ms reward period), and during presentation of the cumulative reward up to that trial in the trial sequence (0-500 ms cumulative reward period). *Perf* neural activations were the relative difference in activity on Δpayoff vs. Δ_0_payoff block RareG trials. Taking the relative block difference allowed non-task related individual EEG differences to cancel out. Relative responses to the RareG door were important for analysis because this door choice resulted in a larger long-term payoff than the other (RareL) door in the Δpayoff block. We applied family-wise error-rate (fwer) corrections to the *Perf* source-space neural correlates to account for multiple comparisons across three frequency bands (theta, alpha, beta) and three time periods (choice, reward, cumulative reward). **Figure 3A** shows the *Perf* source-space neural correlates found to be significant in this analysis; all neural activations inversely related to *Perf*.

**Figure 3:**
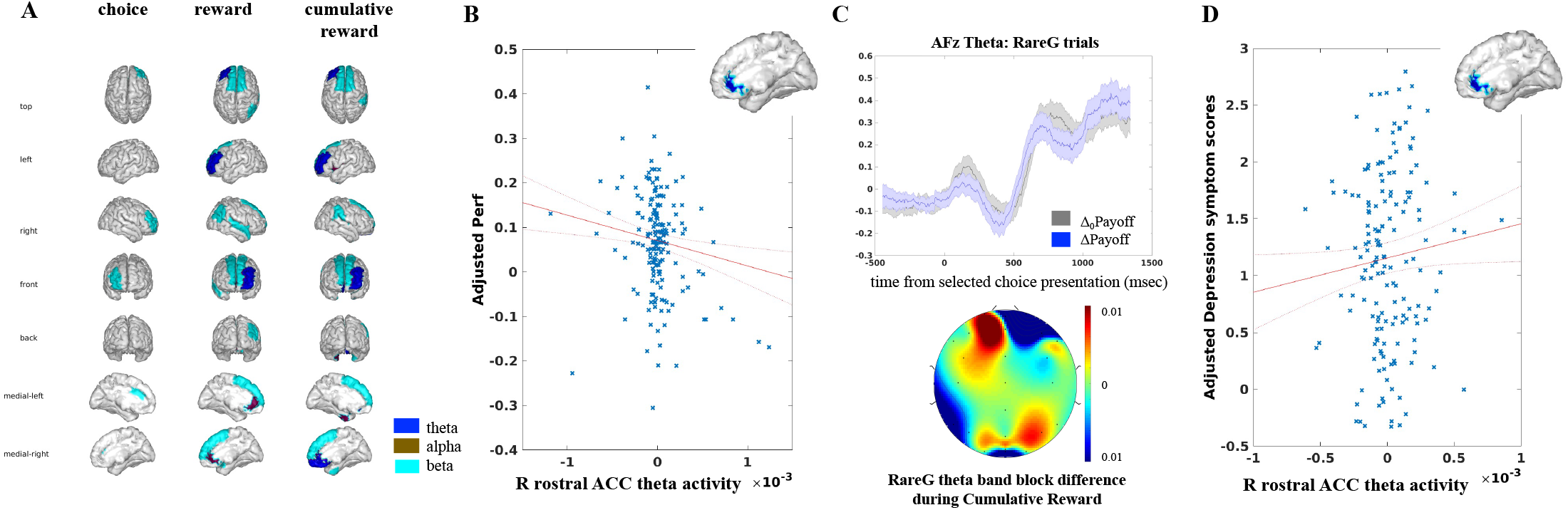
Neural correlates of payoff-based decision making in humans. **(A)** Independent neural correlates in theta (blue), alpha (brown), and beta (cyan) bands found to be inversely related to *Perf* in the choice selection, reward and cumulative reward epochs (p<0.0055). **(B)** Only right rostral anterior cingulate cortex (rACC) theta activity in the cumulative reward period uniquely relates to *Perf* after controlling for *Bias* and time scale of reward prediction. **(C)** Corresponding theta band temporal activity on RareG trials at mid-frontal scalp channel AFz showing differences between payoff blocks (top), and scalp topography of RareG theta band trial difference across Δpayoff and Δ_0_payoff blocks in the significant cumulative reward period (bottom). **(D)** *Perf* related rACC theta positively predicts depressed mood symptoms. The scatter plots in (B) and (D) are presented on an adjusted axis as obtained from the multivariate robust regression models; the x-axes for these plots are 10^−3^ source activity units.

We further accounted for the multiple independently significant cortical ROI predictors (Figure 3A) within a unified multivariate model for *Perf* that also included the significant *Bias* and RL model parameter γ as covariates. This overall multivariate model for *Perf* was significant (R^2^=0.69, df=126, p<0.0001). The results of this model showed alpha activity in the bilateral rostral anterior cingulate cortex (rACC) during the reward period (left: β=-394.32 ± 125.56, t(177)=-3.14, p=0.002; right: β=-634.89 ± 220.51, t(176)=-2.88, p=0.004) and theta activity in the right rACC during the cumulative reward period (β=-56.98 ± 17.93, t(177)=-3.17, p=0.002) as the most significant predictors of payoff-based decisions; activity in the selected choice presentation period did not survive the multivariate model. We then investigated whether these specific theta/alpha rACC activations also related to gain frequency *Bias*, while controlling for the *Perf* payoff. The bilateral alpha band rACC activations were significantly related to *Bias* (left: β=-514.67±205.05, t(178)=-2.51, p=0.01; right: β=-886.15±348.74, t(179)=-2.54, p=0.01) but not right rACC theta (p>0.05). Thus, right rACC theta activity during the cumulative reward period was the distinct neural correlate of reward payoff **(Figure 3B)**; corresponding scalp theta activity and topography are shown in **Figure 3C**.

Additionally, we checked whether the distinct Perf correlate of right rACC theta activation during the cumulative reward period showed any interactions with age and gender, but no significant interactions were found (p>0.1).

### Distinct neural correlates of reward maximization in right rACC predict depressed mood

Finally, we investigated whether the neural correlates of payoff decisions are relevant to subjective mental health by modeling anxiety, depression, inattention and hyperactivity self-reported mood symptoms scores as dependent variables in robust multivariate regression models. All demographic variables (age, gender, race, ethnicity, SES), task performance variables of *Perf* and *Bias*, the relevant RL model parameter γ, as well as the reward-processing neural correlates (bilateral rostral ACC alpha; right rostral ACC theta) were included in each model as independent predictors. Models were fwer-corrected for multiple comparisons.

Only the models for anxiety (R^2^=0.24, p=0.0002) and depression (R^2^=0.21, p=0.002) were significant after multiple comparisons correction. Amongst demographics, age negatively predicted both anxiety (β=-0.02±0.004, t(152)=-4.95, p<0.0001) and depression (β=-0.01±0.004, t(152)=-3.42, p=0.0007); no other demographics, RL model parameter γ, Perf or Bias were significant in these models. Notably, neural correlates of payoff performance, specifically right rACC theta during the cumulative reward period positively predicted depressed mood (β=301.41±150.34, t(152)=2.00, p=0.046, **Figure 3D**); payoff neural correlates did not predict anxiety.

## Discussion

Reinforcement learning models suggest healthy human choices ideally tend to maximize long-term beneficial outcomes (Balasubramani, Chakravarthy, Ali, et al., 2015a; Balasubramani et al., 2014; Balasubramani, Moreno-Bote, et al., 2018; Balasubramani & Chakravarthy, 2019; A. Gupta et al., 2013). However, many existing neuropsychological measures of decision-making that optimize for long-term payoffs don’t reliably estimate the participant’s ability to integrate rewards and make foresighted decisions, and instead suffer from biased predispositions (Furl, 2010; Gansler et al., 2011; Napoli & Fum, 2010; Stocco et al., 2009); this tendency to choose lesser local minima based gain over long-term gain is also referred to as melioration (Sims et al., 2013). In our paradigm, we study gain frequency driven bias separate from payoff-based decision-making by introducing a Δ_0_payoff block with no payoff difference between choice options wherein decisions are purely based on gain frequency (Lin et al., 2009). Comparing choices within the Δpayoff experimental block, designed to have similar reward distribution structure as the Δ_0_payoff block but differing only on the long-term outcome between options, allows measurement of long-term payoff maximization strategy. Therefore, our study design is able to distinguish reward maximization from melioration and further leverages these measures to inform mental health behaviors.

The behavioral outcomes of our experiment varied based on individual subject characteristics. We found that payoff-based performance was significantly related to individual bias for observed frequency of gains; this is in line with prior studies of decision-making but wherein gain frequency decisions are often conflated with expected value (Bechara et al., 1997; Lin et al., 2009). We further modeled subjective differences in reinforcement learning based decision making and by using RL modeling framework, we extracted subjective sensitivity to risks (α) and time scale of reward prediction (γ) to explain each subject’s behavior. The parsimonious RL models suggested that the observed behavior is preferably explained by the differences in the extent of reward prediction over time (γ) between individuals.

Uniquely, we then investigated the neural correlates of payoff performance while accounting for individual differences in gain frequency bias and extent of reward prediction. Earlier studies have suggested complex dynamics mediated by dopamine and serotonin neuromodulation in the cortico-basal ganglia circuit to underlie reward and risk based decision making (Balasubramani, Chakravarthy, Ali, et al., 2015b; Balasubramani, Chakravarthy, et al., 2018; Balasubramani et al., 2014; Chakravarthy et al., 2018; Doya, 2002; Schultz, 2000; Sutton & Barto, 1998; Tobler et al., 2009). In our study, we focused on three different time periods of the task, the first associated with processing of the selected choice, the ensuing reward presentation period and the cumulative reward period to understand how neural dynamics in these periods affect payoff-based performance. The selected choice period captures the processing associated with presentation of the chosen door after the actual decision period. We did not analyze the actual decision period since two different choice options are shown on the screen during this time and the signal associated with every choice option was difficult to explicitly assess. These analyses showed distinct neural correlates of payoff based performance in the core frontal executive region of the right rACC during the cumulative reward period. Relatedly, most prior studies on probabilistic reward processing have suggested this medial prefrontal cortex (mPFC) region to be critical for mediating decision performance (Balasubramani & Hayden, 2018; Bechara et al., 1997, 2000; Brevers et al., 2013; Kennerley et al., 2011; Moccia et al., 2017).

More specifically, analyses showed theta activity in right rACC negatively associated with payoff-based performance. This finding is aligned with prior evidence for reward-based theta processing and its widely studied relationship to long-term risk or uncertainty (Cavanagh et al., 2010; Christie & Tata, 2009; Zavala et al., 2018). This may be one reason why we observe a negative relationship between cumulative reward period rACC theta and effective payoff, *Perf*, because long-term payoff magnitude has been shown to inversely relate to uncertainty but positively to choice utility (Behrens et al., 2007; Krain et al., 2006; Paulus & Frank, 2006; Preuschoff et al., 2006). Notably, reward period bilateral rACC alpha also related to payoff maximization, but this was not a distinct correlate of payoff as it also significantly explained gain frequency bias or melioration in our task.

Translational neuroscience studies show that reward based decision processing deficits are found in depression and in attention disorders, leading to difficulty in reward integration and foresighted choice-behaviors (Balasubramani, Chakravarthy, Ali, et al., 2015b; Balasubramani & Chakravarthy, 2019; Gradin et al., 2011; Groen et al., 2013; Miller & Seligman, 1973; Silvetti et al., 2013; Ziegler et al., 2016). Such individuals then focus on the immediate reward outcome in the short-term, characterized by a prolonged attenuation of temporal discounting of rewards (Eshel & Roiser, 2010; Pizzagalli et al., 2005). Interestingly, rACC theta activity in our paradigm during the cumulative reward period, which negatively correlated with payoff performance, was a positive predictor of depressed mood. This result is aligned with prior research using fMRI showing that the volume and activation patterns of rACC correlates with depressed mood (Boes et al., 2008; Yoshimura et al., 2010). Related research also suggests that rACC theta activity is a significant baseline marker for depression treatment outcomes (Pizzagalli et al., 2018). These results are still limited by our study in healthy adults and need to be replicated in clinical populations, and with greater channel density electrophysiology recordings.

Altogether, our study presents the importance of controlling for melioration biases for immediate reward frequency and individual differences in learning while assessing advantageous, i.e. foresighted, decision-making ability in humans. Our findings of payoff-relevant right rostral ACC theta activity in the cumulative reward feedback period, could be important for clinical translational application, particularly for depression, suggesting a plausible neural target for interventions that engage reward processing.

## Supplementary Data

**Supplementary Table 1.**
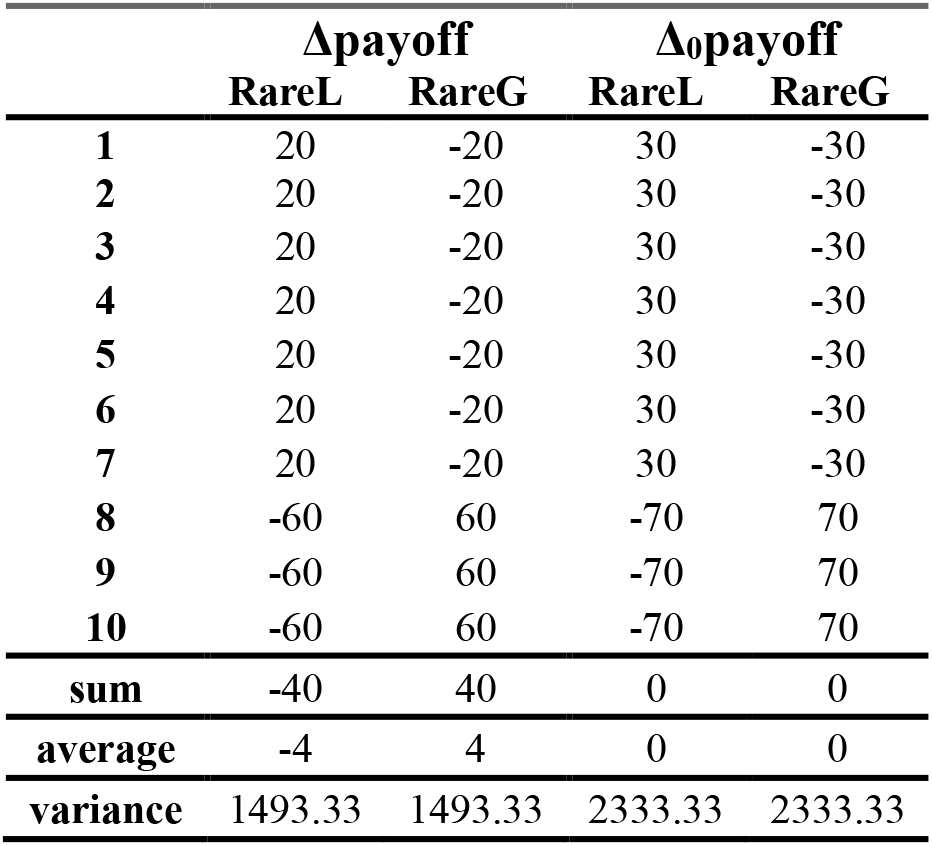
Reward distributions for the door choices in the Δpayoff and Δ_0_payoff blocks. The two door choices in either block were RareG (rare gains and frequent losses) and RareL (rare losses and frequent gains). Payoff (expected value) for RareG and RareL were the same in the Δ_0_payoff block and greater for RareG relative to RareL in the Δpayoff block. RareG and RareL distributions had the same sum, average and variance in the Δ_0_payoff block, and different sum and averages but same variance in the Δpayoff block.

## Acknowledgements

This work was supported by University of California San Diego (UCSD) lab start-up funds (DR, JM), the Interdisciplinary Research Fellowship in NeuroAIDS (PB: R25MH081482), the Brain & Behavior Research Fund (PB), the Kavli Foundation (PB, JM), and the Sanford Institute for Empathy and Compassion (JM, PB). We thank Alankar Misra for software development of the *BrainE* software and several UCSD undergraduate students who assisted with data collection. The *BrainE* software is copyrighted for commercial use (Regents of the University of California Copyright #SD2018-816) and free for research and educational purposes. We thank Sabyasachi Shivkumar and Vignesh Muralidharan for their helpful feedback on the study analysis.

## Data Availability

A part of the dataset with 96 of the 186 participants data used in this study is available on the open-access repository link: 10.5281/zenodo.4088951

